# ATR function is indispensable to allow proper mammalian follicle development

**DOI:** 10.1101/471698

**Authors:** Sarai Pacheco, Montserrat Garcia-Caldés, Ignasi Roig

**Affiliations:** Genome Integrity and Instability Group, Institut de Biotecnologia i Biomedicina, Universitat Autònoma de Barcelona, Cerdanyola del Vallès, Spain; Department of Cell Biology, Physiology and Immunology, Universitat Autònoma de Barcelona, Cerdanyola del Vallès, Spain; MRC Clinical Sciences Centre, Faculty of Medicine, Imperial College London, Du Cane Road, London W12 0NN, UK

**Keywords:** Folliculogenesis, ATR, oogenesis, replication, apoptosis

## Abstract

Mammalian female fertility relies on the proper development of follicles. Right after birth in mouse, oocytes associate with somatic ovarian cells to form follicles. These follicles grow during adult lifetime to produce viable gametes. In this study, we analyzed the role of the ATM and rad3-related (ATR) kinase in mouse oogenesis and folliculogenesis using a hypomorphic mutation of the *Atr* gene (Murga et al., 2009). Female mice homozygote for this allele have been reported to be sterile. Our data show that female meiotic prophase is not grossly altered when ATR levels are reduced. However, follicle development is majorly compromised since *Atr* mutant ovaries present a decrease of growing follicles. Comprehensive analysis of follicular cell death and proliferation suggest that wild-type levels of ATR are required to achieve optimal follicular development. Altogether, these findings suggest that reduced ATR expression causes sterility due to defects in follicular progression rather than in meiotic recombination. We discuss the implication of these findings for the use of ATR inhibitors as anti-cancer drugs and its possible side effects on female fertility.

## Introduction

Gametogenesis is the process that generates mature haploid cells or gametes, that can participate in fertilization. Mammalian gametogenesis is characterized by a sexual dimorphism that reflects in different aspects of the process (Morelli and Cohen, 2005). For instance, while males produce sperm during most of their adult life, females are born with a limited amount of germ cells, which sets the extension of the fertile life of the animal. Mammalian female gametogenesis, or oogenesis, initiates during fetal time when oogonia differentiate into oocytes and enter into meiotic prophase. After completing, homologous chromosome synapsis and meiotic recombination, oocytes arrest at the end of the meiotic prophase, called dictyate. Then, oocytes surround by somatic ovarian stromal cells to form primordial follicles, in which oocytes enter a quiescent state that may last decades depending on the species. It is widely accepted that these primordial follicles represent the pool of gametes each female will have during their adult life. Folliculogenesis initiates, once these dormant primordial follicles are recruited to start the growing process. Folliculogenesis involves a remarkable increase in the size of the oocyte accompanied by a substantial proliferation of granulosa cells that surround the oocyte, increasing the total volume of the follicle more than 100-folds (van den Hurk and Zhao, 2005). Intercellular communication between the oocyte and the follicular cells is crucial to allow proper oocyte maturation (Li and Albertini, 2013; Sánchez and Smitz, 2012). Folliculogenesis ends with the formation of a preovulatory follicle which contains an oocyte capable of resuming meiosis (Li and Albertini, 2013).

The fact that mammalian females are born with the pool of oocytes they will use in their lifetime, represents a challenge since these cells are exposed to several environmental aggressions during the life of the individual that may ultimately harm them. Among others, most cancer therapy treatments are known to produce infertility as a side-effect due to the cessation of follicle pool (Woodard and Bolcun-Filas, 2016). Thus, many efforts are being invested in finding ways to preserve fertility for cancer patients (Woodard and Bolcun-Filas, 2016). Some of these efforts involve the use of more selective drugs against cancer cells that do not damage other cell types, like germ cells. One example is ATR inhibitors, which exploit the fact many tumor cells suffer oncogene-induced replication stress (Karnitz and Zou, 2015; Murga et al., 2011). ATR (ATM and rad3-related) kinase is a master regulator of DNA damage response in mammalian cells (Blackford and Jackson, 2017). ATR is activated by the presence of long stretches of RPA-coated single stranded DNA (Zou and Elledge, 2003). These mainly arise from stalled replication forks, but can also be originated by the processing of several other forms of DNA damage (Blackford and Jackson, 2017). The activation of ATR leads to a delay in cell cycle progression, the prevention of replication forks collapse and the repair of DNA damage (Blackford and Jackson, 2017; Nam and Cortez, 2011). Thus, those tumor cells that present replication stress depend on ATR to stabilize the replication forks and delay enter into mitosis to resolve these problems and survive. In the absence of ATR, or a functional checkpoint, these cells are unable to repair these lesions and suffer replication catastrophe and ultimately die (Toledo et al., 2017).

The study of ATR functions in mammalian gametogenesis has been difficult because ATR is essential for embryo development (Brown and Baltimore, 2000). Nonetheless, the generation of a mouse model carrying a hypomorphic mutation responsible for the Seckel syndrome in humans, that reduces the expression of ATR, but it is still compatible with life, has allowed a better understanding of ATR functions *in vivo* (Murga et al., 2009). Seckel mice show high levels of replicative stress associated with embryonic cell proliferation. In adult tissues, Seckel mice showed signs of premature aging, possibly due to the depletion of the stem cell pool (Murga et al., 2009). Remarkably, female Seckel mice presented a severe dwarfism and were reported to be sterile, since no viable oocytes were obtained after hormone-induced superovulation. This study suggested that wild-type levels of ATR expression are required for the ovulation of fertilizable oocytes. The authors proposed that this might be due to defects in the completion of meiotic recombination in Seckel oocytes.

In this study, we have analyzed oogenesis progression in Seckel mouse in order to characterize the role of ATR in female fertility. Our data shows that contrary to our expectations, Seckel mouse oocytes complete homologous chromosome synapsis and meiotic recombination without noticeable defects. However, our detailed analysis of folliculogenesis progression shows that adult Seckel mouse ovaries present a reduction of growing follicles. Moreover, these growing follicles are smaller than those found in control ovaries and have more TUNEL-positive cells and less proliferating cells than control ones. Altogether, our results suggest that ATR is required to permit proper follicular cells proliferation. These data imply that the use of ATR inhibitors may alter follicular cell proliferation leading to transient infertility.

## Results

### Meiotic prophase progression is not altered in Seckel mouse oocytes

As mentioned earlier, female Seckel mice have been proposed to have meiotic recombination defects (Murga et al., 2009), since we have previously demonstrated that ATR function is required to complete meiotic recombination in spermatocytes (Pacheco et al., 2018; Widger et al., 2018), we studied meiotic prophase progression in Seckel mouse oocytes.

Firstly, we immunostained control and mutant oocytes against proteins of the lateral and central element of the synaptonemal complex (SYCP3 and SYCP1 respectively) to study homologous chromosome synapsis. Chromosome synapsis progressed properly through all prophase stages in Seckel mouse oocytes, indicating that SC formation between homologous chromosomes was comparable in control and mutant oocytes (Fig 1A). This finding suggested that wild-type levels of ATR are not a requirement to achieve a complete homologous chromosome synapsis in mouse oocytes.

**Figure 1.**
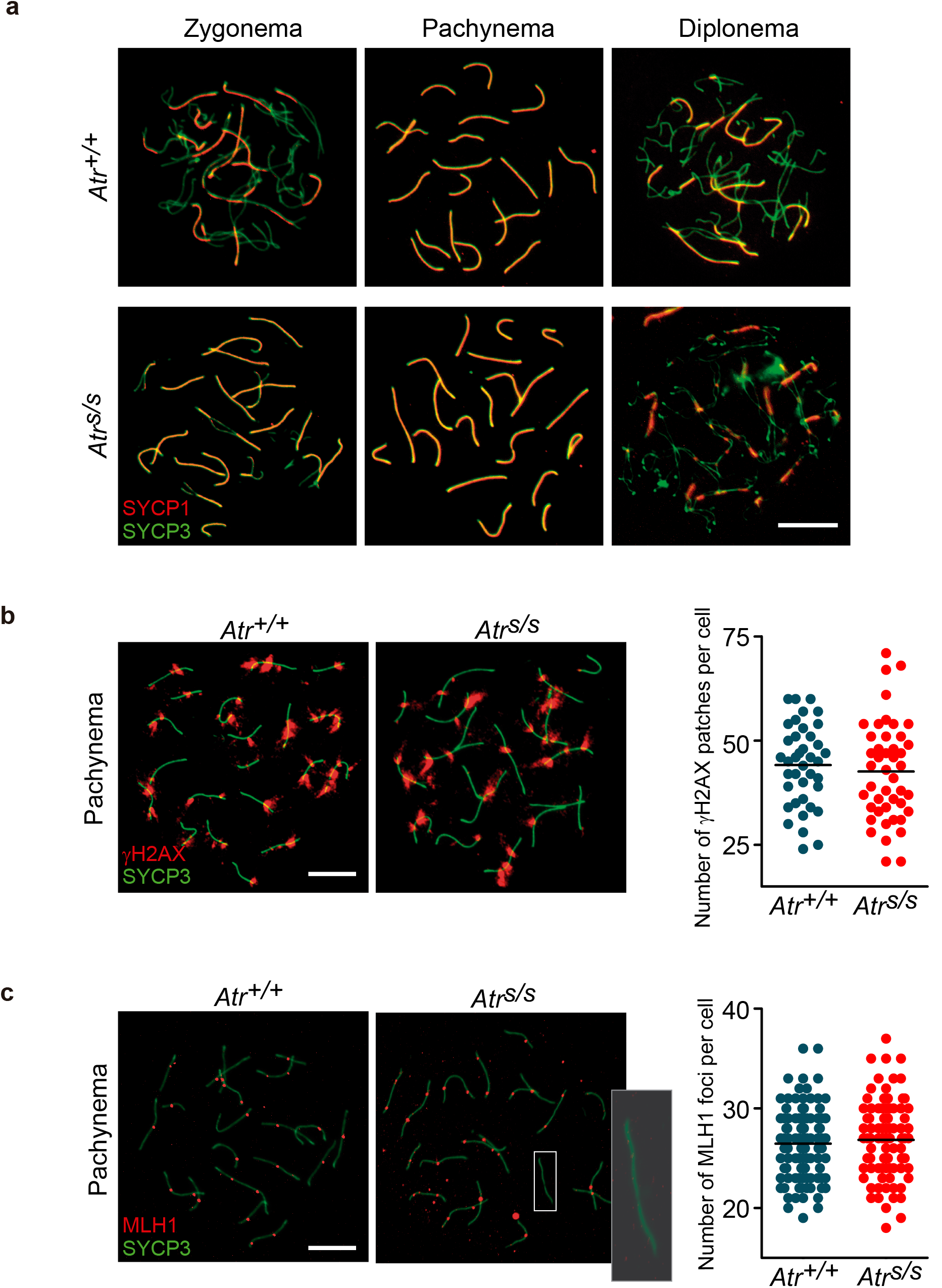
Seckel mouse oocytes exhibit proper meiotic prophase. (**a**) Representative images from *Atr^+/+^* and *Atr^S/S^* oocytes immunostained against SYCP3 and SYCP1 proteins to follow synaptonemal complex formation. Scale bar represents 10 μm and applies to all images. (**b**) Left panels, representative images of oocytes stained against γH2AX and SYCP3. Scale bar represents 10 μm and applies to both images. Right panel, quantification of total γH2AX patches at pachytene stage. Each point represents the number of patches found for a single cell. Horizontal lines denote the mean. (**c**) Left panels, representative images of pachytene oocytes stained against MLH1 and SYCP3. Scale bar represents 10 μm and applies to both images. Right panel, quantification of total MLH1 foci at pachytene stage. Each point represents the number of focus count for a single cell. Horizontal lines denote the mean

Furthermore, to examine how the reduction of ATR levels might affect double strand break (DSB) repair progression, we performed a global analysis of meiotic recombination by immunostaining γH2AX protein on oocyte spreads. In mammalian oocytes, γH2AX signals SPO11-generated DSBs showing discrete staining patches along chromosomes axes at pachynema (Roig et al., 2004). We found no differences regarding the total number of γH2AX patches found in mutant versus control oocytes at pachynema (44.1 ± 1.6 in *Atr^+/+^* (mean ± SD, N=39), 42.6 ± 1.8 in *Atr^s/s^* (N=45), p=0.5322, t test, Fig. 1B). These results suggest that, in contrast with what we observed in mutant males, in Seckel mouse oocytes, meiotic recombination progression is not obviously disturbed.

We also studied one of the outcomes of meiotic recombination, crossover (CO) formation, in Seckel mouse oocytes. DSB repair by meiotic recombination can lead to the exchange of genetic material between the homologous chromosomes, known as CO, or just the use of one homologous chromosome as the template to repair the other, which is called non-crossover. Because COs are crucial to maintaining homologous chromosomes together until metaphase I, CO formation is tightly regulated in order to ensure that at least one CO is formed per each homologous pair (Cole et al., 2012). We immunostained wild-type and mutant oocytes against MLH1, which marks most COs in mammals (Edelmann et al., 1996)(Fig. 1C). The total number of MLH1 foci present in pachytene oocytes was identical between control and mutant mice (26.5 ± 0.3 in *Atr^+/+^* (mean ± SD, N=107); 26.8 ± 0.4 in *Atr^s/s^* (N=95), p=0.4826, Mann Whitney test). Interestingly, as it occurs in spermatocytes (Pacheco et al., 2018), Seckel mouse oocytes presented a significant increase in the number of bivalents without an MLH1 foci at pachynema (2.1% of the bivalents in *Atr^+/+^* (N=2140); 3.9% of the bivalents in *Atr^s/s^* (N=1900), p=0.0010, Fisher’s exact test, inset in Fig. 1C). This finding could be explained if either Seckel mouse oocytes fail to form the obligate CO in an homologous pair or if CO formation may take longer in *Atr* mutant mice due to a mildly impaired meiotic recombination (Pacheco et al., 2018). To test these hypotheses, and in light of MLH1 foci in females persisting longer at COs sites (Baker et al., 1996), we analyzed the number of bivalents lacking MLH1 foci present at diplonema in control and mutant oocytes. We did not find statistically significant differences between both genotypes at diplonema (1.5% of the bivalents in *Atr^+/+^*(N=260); 1.1% of the bivalents in *Atr^s/s^* (N=280), p=0.7164, Fisher’s exact test). These data suggest that the results observed at pachynema may represent a delay in CO formation, rather than an inability to form COs. Together, all these findings support the idea that ATR is required to achieve a proper timely meiotic recombination, even though; meiotic prophase is not grossly altered in female Seckel mice.

### Folliculogenesis progression is altered in adult Seckel mouse ovaries

Since meiotic prophase progression did not seem severely compromised in Seckel female mice, we studied the following stages of oogenesis. In mammals, female fertility depends upon the development of ovarian follicles (Sánchez and Smitz, 2012). After birth, arrested oocytes become surrounded by a single layer of flattened somatic cells to form primordial follicles (Fig. 2A). Primordial follicles rest within the ovary in a quiescent stage until they are recruited to initiate follicle growth, or folliculogenesis. Initiation of folliculogenesis is characterized by the formation of the primary follicles, which contain cubic follicular cells surrounding the oocyte. Folliculogenesis results in an increase in oocyte size. At the same time, follicular cells proliferate mitotically forming a multilayered, or secondary, follicle. Eventually, a cavity appears among follicular cells (antrum) as the follicles develop. The antrum grows until it occupies most of the follicular volume right before ovulation occurs (Sánchez and Smitz, 2012).

**Figure 2.**
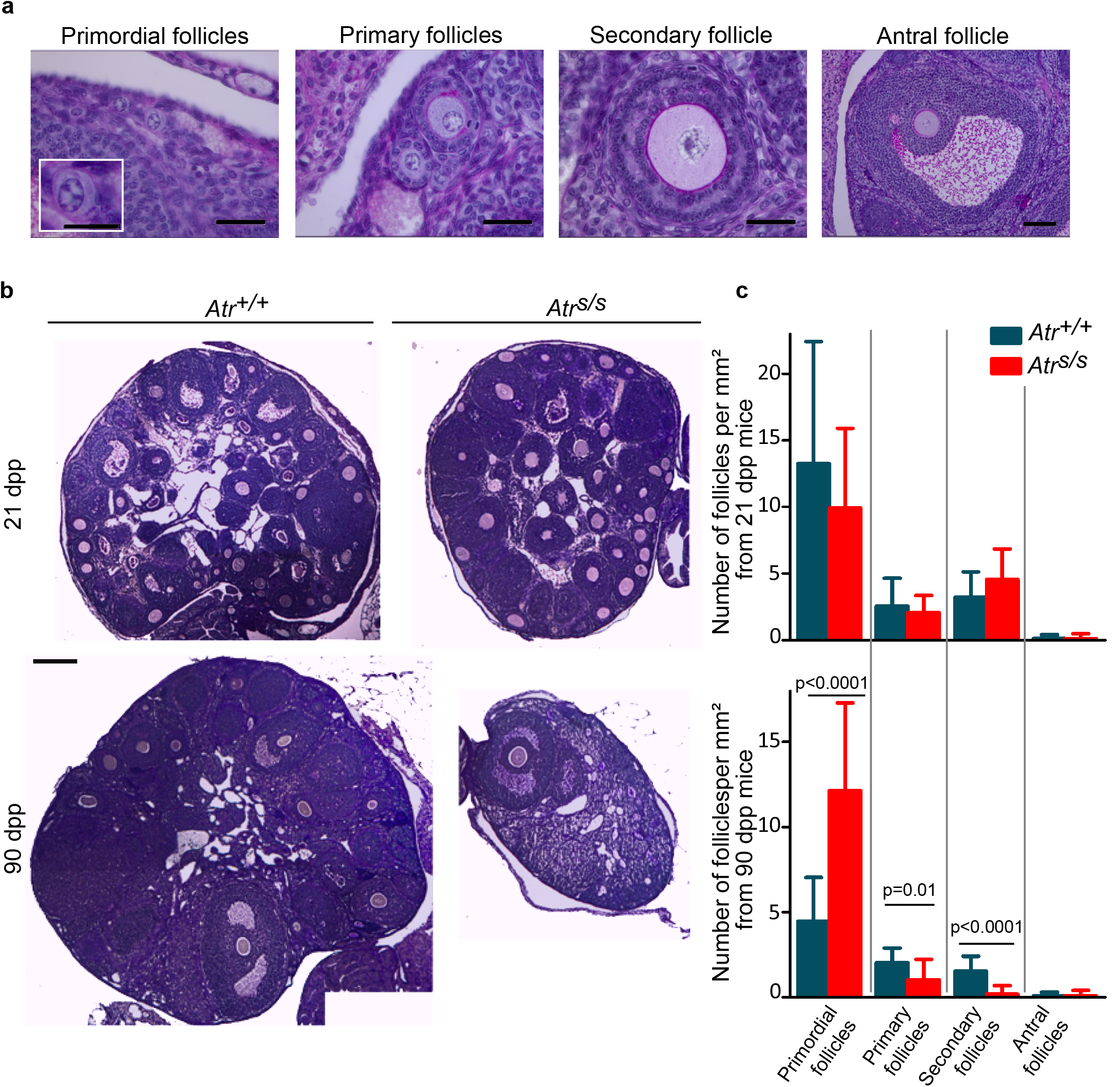
Adult Seckel mouse ovaries present an increased number of primordial follicles, accompanied by a reduction of growing follicles. (**a**) Illustrative images of the different folliculogenesis stages used to classify follicles. Scale bars represent 20 μm, except for the antral follicle that represents 50 μm. (**b**) Histological ovarian sections stained with PAS-Hematoxylin from 21 and 90 dpp mice from *Atr^+/+^* and *Atr^S/S^* ovaries exhibiting all follicular stages. Scale bar represents 200 μm and applies to all images. (**c**) Quantification of the number of follicles present at different stages in eight alternate ovarian sections from two ovaries of the indicated age and genotype. Values are expressed as the average number of follicles present in each section relative to the area of those sections. Columns and lines indicate the mean and standard deviation (SD). Statistically significant differences are indicated with the p values obtained from t tests.

Hence, we counted and classified the follicles from histological sections from control and mutant, prepubertal and adult ovaries into these four different categories according to their morphology (Fig. 2A). As mentioned earlier, since Seckel mice exhibit several dwarfism (Murga et al., 2009), in order to account for the effect of a smaller ovary in this analysis, we measured the total area of each ovarian section and expressed the values as the number of follicles per square millimeter of the ovarian section. Histological analysis revealed that both genotypes contained all follicular stages, in prepubertal as well as in adult ovaries (Fig. 2B), indicating that follicles in Seckel mice have the capacity to progress to pre-ovulatory stages. Nevertheless, although folliculogenesis progression was not affected in prepubertal Seckel ovaries (Fig. 2C, see Table 1), we found a three-fold increase in the number of primordial follicles (p<0.0001, t test) and a significant reduction in primary (p=0.0111, t test) and secondary follicles in adult Seckel ovaries (p<0.0001, t test). Interestingly, similar results were obtained when these numbers were expressed based on a percentage of follicles observed per section, without normalizing per different ovary size (Fig. S1). These data indicate that adult Seckel mice may present defects in primordial follicle recruitment, as well as, in follicular development as suggested by the reduction of growing follicles. Moreover, we also found that Seckel mouse ovaries contained an increased proportion of abnormal follicles. These follicles presented very large oocytes, suggesting they had initiated oocyte growth, but only one layer of flattened or cuboidal granulosa cells (0.83 ± 0.18 follicles/mm^2^ in *Atr^+/+^* (mean ± SD, N=16); 2.1 ± 0.”0 in *Atr^s/s^* (N=16), p=0.0082, Fig. S2). These results suggest that reduction of ATR expression in the ovary alters the correct development of mammalian follicles.

**Table 1.** Average number of follicles per square millimeter present at different stages of folliculogenesis at the indicated age and genotype.

|  |  | Primordial<br>follicles/mm<sup>2</sup> | Primary<br>follicles/mm<sup>2</sup> | Secondary<br>follicles/mm<sup>2</sup> | Antral<br>follicles/mm<sup>2</sup> | N |
| --- | --- | --- | --- | --- | --- | --- |
| 21 dpp<br>ovaries | *Atr*<sup>+/+</sup> | 13.3 ± 9.1 | 2.6 ± 2.1 | 3.2 ± 1.9 | 0.1 ± 0.2 | 16 |
|  | *Atr*<sup>s/s</sup> | 9.9 ± 6 | 2.1 ± 1.3 | 4.5 ± 2.3 | 0.1 ± 0.4 | 16 |
| 90 dpp<br>ovaries | *Atr*<sup>+/+</sup> | 4.6 ± 2.6 | 2.0 ± 0.9 | 1.5 ± 0.9 | 0.09 ± 0.2 | 16 |
|  | *Atr*<sup>s/s</sup> | 12.1 ± 5.2 | 1 ± 1.2 | 0.2 ± 0.5 | 0.08 ± 0.3 | 16 |

Mean ± standard deviation and total number of sections analyzed (N) are shown.

### Follicular cell proliferation is compromised in Seckel mouse ovaries

The correct proliferation of follicular cells is necessary not only for follicular growth and development but also for the establishment of an appropriate environment for oocyte maturation (Li and Albertini, 2013). To study whether follicular growth was compromised in Seckel mouse ovaries, we indirectly evaluated follicular cell proliferation. To do so, we measured the total area of a follicle and the area of its oocyte in control and Seckel mouse follicles. Then, we expressed this data as a relative area occupied by the oocyte in a particular follicle (Fig. S3 and Table S1). In control mice, the relative area of an oocyte decreased throughout folliculogenesis progression, indicating an increase in the area occupied by follicular cells, as a result of the cell proliferation occurring in these stages. Interestingly, although in Seckel mice the relative oocyte area also tended to decrease throughout folliculogenesis, the area occupied by oocytes in early growing follicles was larger than in control mice. Thus, while examination of primordial follicles did not show statistically significant differences between control and mutant follicles (p = 0.1474, t test, see Supplementary Table 1), Seckel ovaries presented larger oocytes enclosed in primary ((p<0.0001, t test), and small secondary follicles (with up to three layers of granulosa cells, p<0.0001, t test). Remarkably, we found indistinguishable relative area between control and Seckel pre- antral secondary follicles (with more than three layers of granulosa cells, p=0.7360, t test) and, antral follicles (p=0.4884, t test). These data suggest that Seckel mouse follicles contain a reduced number of granulosa cells surrounding the oocyte as compared to control mice.

Coordination of cell populations are strictly controlled by the balance between cell proliferation and cell loss. Thus, successful follicle development depends in part on follicular cell proliferation and prevention of apoptosis (Liu, 2007). Since regular follicular development is accompanied by high proliferation rate of follicular cells we inquired whether the reduced ATR expression observed in the ovary might be altering cell proliferation and programmed cell death during folliculogenesis. Since we only observed developmental problems in primary and small secondary follicles, we focused the following analyses on these two categories of follicles.

Firstly, to evaluate granulosa cell proliferation, we used an EdU incorporation assay to detect replicating follicular cells on cryosections from prepubertal and adult ovaries (Fig. 3). We analyzed the number of EdU positive cells relative to the total number of granulosa cells present in a particular follicle. Importantly, the number of granulosa cells counted in *Atr* mutant follicles tended to be lower than the observed in control follicles (Table 2), consistent with what we had indirectly determined before. EdU analyses revealed that follicular cell proliferation rate in Seckel mouse primary follicles tended to be lower than the observed in control follicles, being statistically significant only in adult mice (p=0.0003, t test). Interestingly, we observed an increased proportion of EdU-positive cells in mutant small secondary follicles, both in prepubertal (p=0.0144, t test) and in adult ovaries (p=0.0008, t test). These findings suggest that Seckel follicles have a reduced granulosa cells proliferation rate, which strengthens as mice age, leading to problems on follicular development and folliculogenesis progression. Nevertheless, due to the increased proliferation rate detected in secondary follicles, we suggest that those follicles that have the ability to progress throughout folliculogenesis present an increased granulosa cell proliferative capacity, which allows them a proper follicular development.

**Table 2.** Average follicular cell proliferation, EdU positive cells and number of follicular cells at the indicated age and genotype.

|  |  |  | Proliferation rate | EdU positive cells | Number of follicular cells | N |
| --- | --- | --- | --- | --- | --- | --- |
| **Prepubertal mice** | Primary follicles | *Atr<sup>+/+</sup>* | 0.16 ± 0.17 | 3.0 ± 3.8 | 15.1 ± 6.7 | 34 |
|  |  | *Atr<sup>S/S</sup>* | 0.1 ± 0.1 | 1.7 ± 2.3 | 11.9 ± 6.2 | 36 |
|  | Secondary follicles | *Atr<sup>+/+</sup>* | 0.2 ± 0.1 | 13.8 ± 8.5 | 70 ± 30.2 | 67 |
|  |  | *Atr<sup>S/S</sup>* | 0.24 ± 0.1 | 13.9 ± 6.6 | 57.6 ± 21.5 | 57 |
| **Adult mice** | Primary follicles | *Atr<sup>+/+</sup>* | 0.1 ± 0.1 | 2.3 ± 3.4 | 18.1 ± 8.3 | 28 |
|  |  | *Atr<sup>S/S</sup>* | 0.03 ± 0.07 | 0.4 ± 0.8 | 10.5 ± 4.5 | 43 |
|  | Secondary follicles | *Atr<sup>+/+</sup>* | 0.10 ± 0.08 | 5.7 ± 5.10 | 62.8 ± 27.1 | 56 |
|  |  | *Atr<sup>S/S</sup>* | 0.17 ± 0.06 | 7.2 ± 3.5 | 43.4 ± 15.8 | 18 |

Mean ± standard deviation and total number of follicles analyzed (N) are shown.

**Figure 3.**
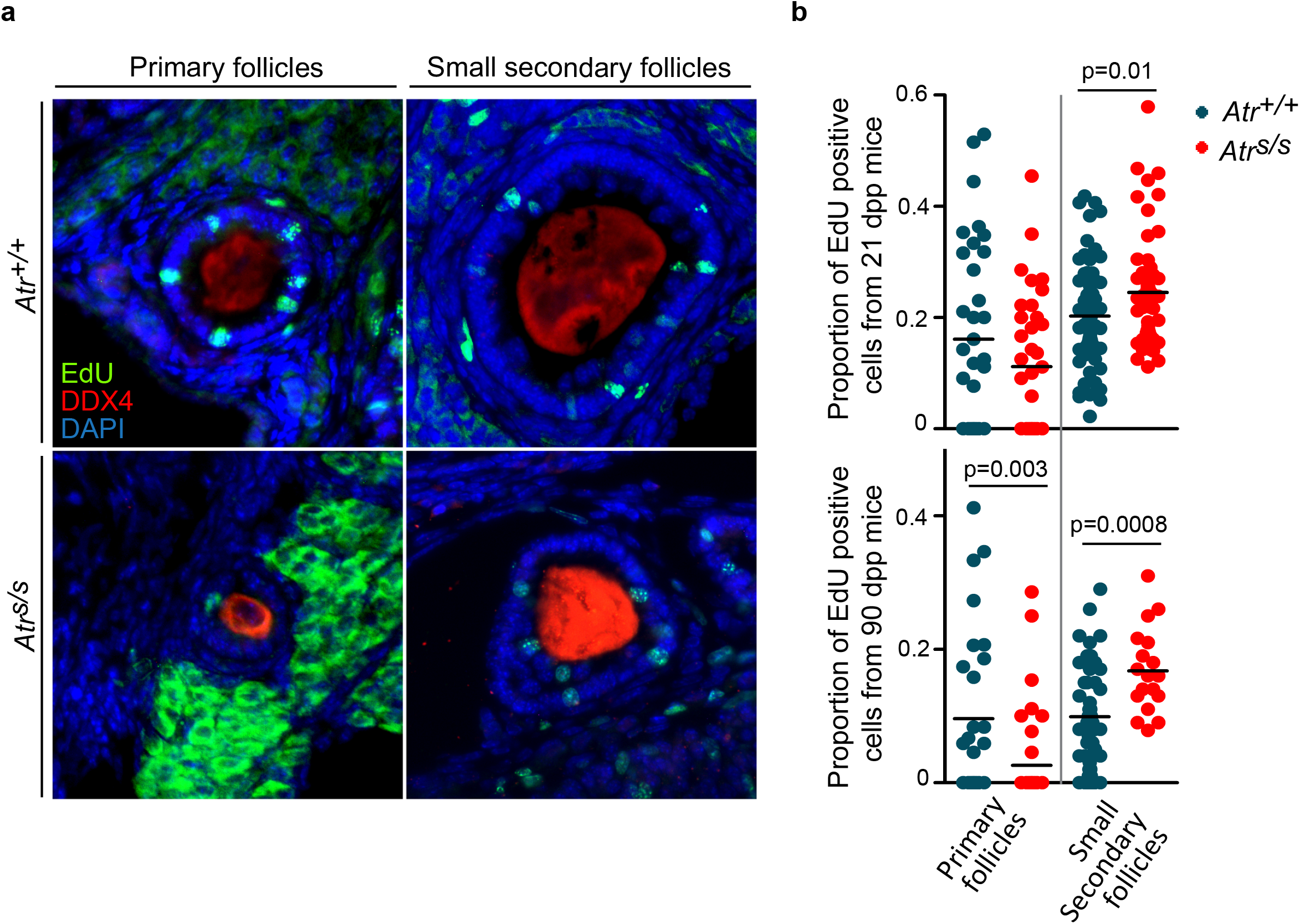
Granulosa cell proliferation rate is affected in Seckel follicles. (**a**) Representative images of primary and small secondary follicles showing EdU positive cells, DDX4 and DAPI from *Atr^+/+^* and *Atr^S/S^* ovaries. Scale bar represents 50 μm and applies to all images. (**b**) Quantification of the proportion of EdU positive cells relative to the total number of granulosa cells present in each follicle. Horizontal line denotes the mean. Adult Seckel females exhibit significantly reduced proportion of EdU positive granulosa cells in primary follicles, but an increased proliferation rate in small secondary follicles.

Next, we studied follicular cell death performing TUNEL assays on histological sections from prepubertal and adult ovaries from control and Seckel mice. We determined the apoptotic follicular cell rate by analyzing the number of TUNEL positive cells relative to the total number of granulosa cells present in a particular follicle (Fig. 4 and Table 3). Seckel mouse follicles tended to present more apoptotic cells than control follicles in all follicular stages. Nevertheless, in this case, while in primary follicles apoptotic cell rate was not statistically different between control and Seckel mouse ovaries, neither in prepubertal (p=0.”8”9, t test), not in adult mice (p=0.5967, t test), small secondary follicles presented a significant increase of follicular cell apoptotic rate (five-fold increase in prepubertal, p<0.0001, t test, and a nine-fold increase in adult mice, p=0.0017, t test). This increment in the apoptotic follicular cells rate present in Seckel ovaries supports the idea that reduced ATR expression in the ovaries causes follicular cell defects.

**Table 3.**
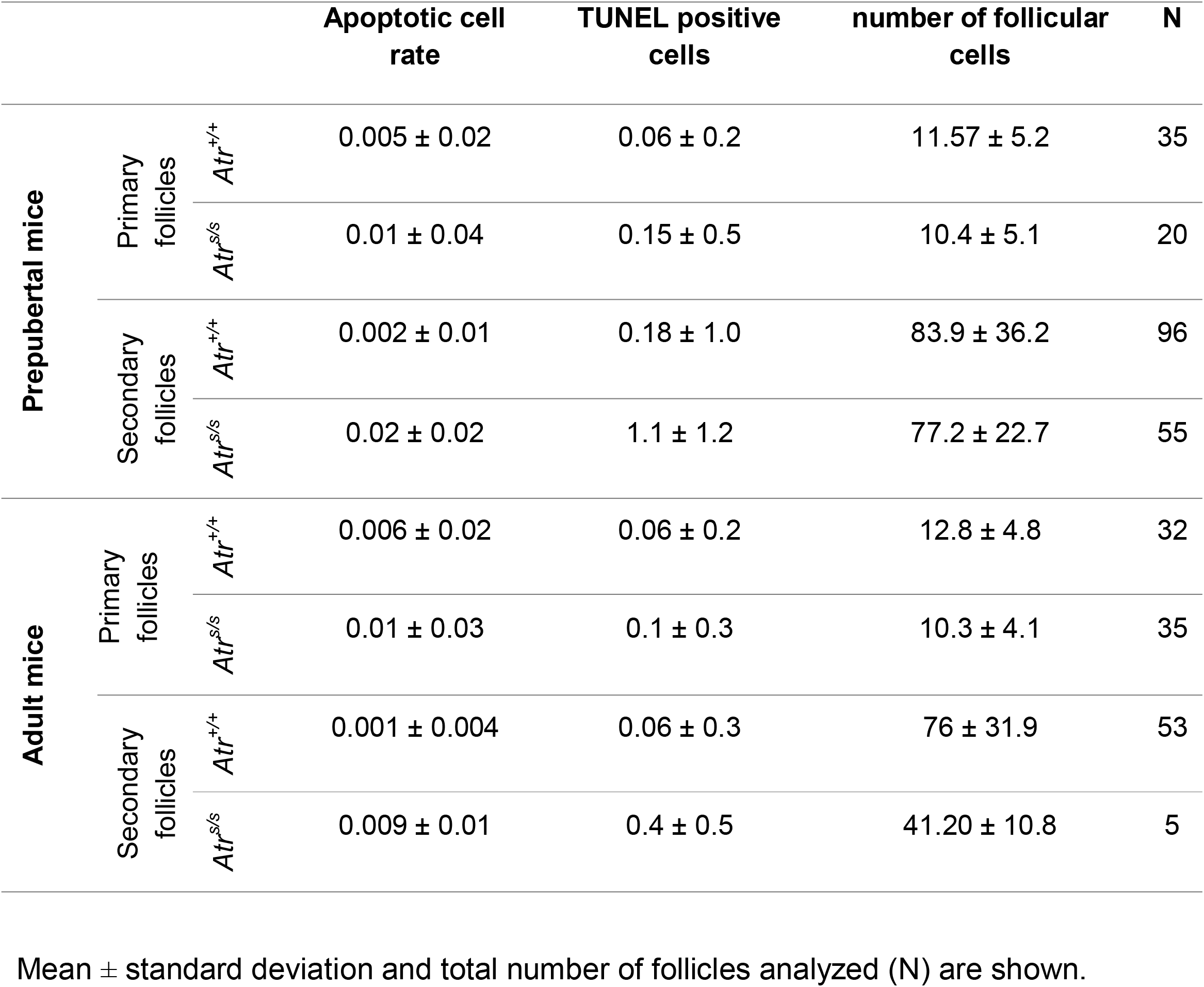
Average apoptotic follicular cell rate, TUNEL positive cells and number of follicular cells at the indicated age and genotype.

**Figure 4.**
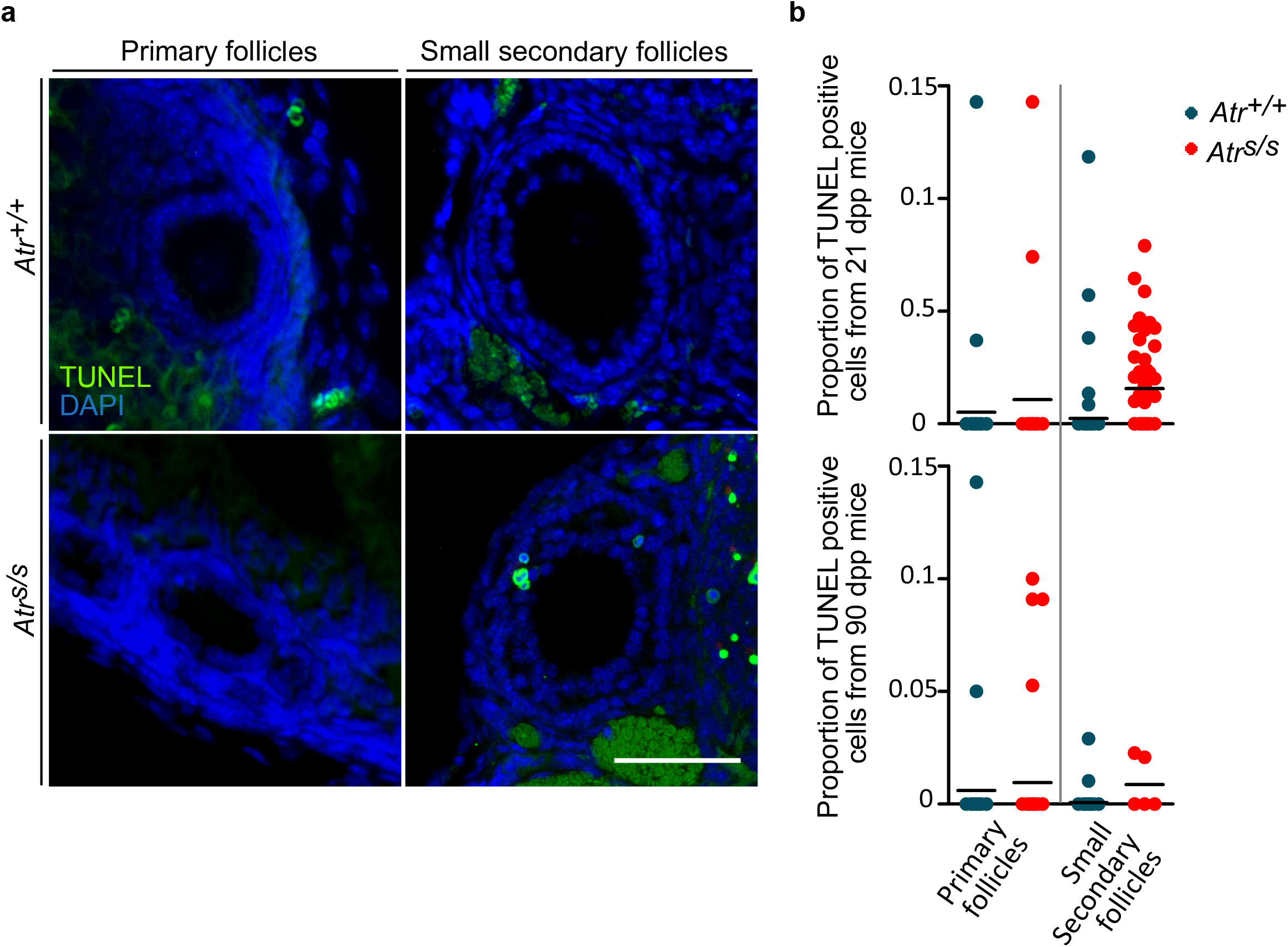
Seckel mouse follicles exhibit an increased proportion of apoptotic granulosa cells. (**a**) Representative images of primary and small secondary follicles showing TUNEL positive cells, and DAPI from *Atr^+/+^* and *Atr^S/S^* ovaries. Scale bar represents 50 μm and applies to all images. (**b**) Proportion of TUNEL positive cells relative to the total number of granulosa cells present in each follicle. Horizontal line denotes the mean.

Hence, these data indicate that in Seckel mouse ovaries, while follicles grow and the number of granulosa cells rises, the number of apoptotic granulosa cells also increases, intensifying the developmental problems that experience these follicles. Consistent with proliferation analysis, these results suggest that only those follicles that achieve a high proliferative capacity can overcome the increased apoptotic rate present in Seckel mouse follicles and thus progress throughout folliculogenesis. Finally, all these findings suggest that ATR is necessary to achieve a successful follicle development and consequently, a correct folliculogenesis progression that enables fertility.

## Discussion

It is well known that coordinated regulation of oocyte and follicle growth is essential for female fertility. Therefore, proficient communication between oocyte and their surrounding follicular cells during folliculogenesis plays a crucial role to develop a healthy oocyte ready for ovulation and fertilization (Li and Albertini, 2013; Sánchez and Smitz, 2012). Here we demonstrated reduced ATR expression in Seckel ovaries originates follicular developmental defects that may have critical consequences in fertility.

Seckel females were reported to be infertile presumably due to meiotic recombination defects (Murga et al., 2009). However, our experimental findings showed that while meiotic prophase appeared to be normal, histological analysis showed unusual disturbances in Seckel mouse ovarian sections that were more pronounced in adult ovaries. We observed a severe reduction of growing follicles in Seckel adult ovaries, despite the presence of few antral follicles. In addition, mutant adult sections presented abundant stroma devoid of follicles, which could be associated to the accelerated aging related to Seckel mutation (Murga et al., 2009).

ATR responds to DNA damage controlling replication origin firing, preventing replication forks collapse, regulating cell cycle progression, preventing premature mitotic entry and promoting DNA repair (Blackford and Jackson, 2017; Sørensen et al., 2005). Deficient ATR function could lead to errors produced by replication fork collapse, accumulating DNA damage and ultimately causing cell death (Cimprich and Cortez, 2008). During follicular development, granulosa cells actively proliferate to generate multilayered antral follicles. Here we showed that granulosa cells from Seckel mouse primary follicles display proliferation defects. We observed that in adult primary follicles the proliferation rate from *Atr* mutant granulosa cells was lower than the one found in control cells. We interpret these results as a consequence of the increased replication stress associated to the follicular development occurring in an ATR-defective scenario. As mentioned earlier, follicular development is associated with an increased proliferation of follicular cells, this may result in replication stress, which leads to mitotic catastrophe in *Atr* mutant cells (Lecona and Fernández-Capetillo, 2014; Toledo et al., 2017). Strikingly, granulosa cells from mutant secondary follicles showed a greater proliferation rate than control cells. We think this is a manifestation of the selection that occurs within the follicle pool at the first stages of folliculogenesis. Only those follicles that are able to overcome the replication stress, and the consequent loss of cells associated to a reduced *Atr* expression may be able to progress to more advanced stages of folliculogenesis. Interestingly, similar selection processes have been proposed to occur in other adult tissues of Seckel mouse (Murga et al., 2009). In our opinion, this explains why Seckel mouse ovaries contain less growing follicles than wild type mice and may be the reason of their infertility.

Our data also demonstrate that adult Seckel ovaries have an impaired recruitment of primordial follicles, given that the pool of dormant follicles observed was threefold higher in mutant mouse ovaries than in control ovaries. This observation suggests that ATR may play a role in the activation of primordial follicles. Primordial follicles are selectively and continuously recruited to progress throughout folliculogenesis in a process mainly regulated by germ cell specific proteins (Pelosi et al., 2015). Nonetheless, it has been demonstrated that some somatic cell signaling factors are also essential to activate primordial follicles (Kerr et al., 2013; Pelosi et al., 2015; Zhang et al., 2014;). Thus, it is not clear from our observations if ATR function may be required in the oocyte, the follicular cells or both in order to recruit primordial follicles.

Interestingly, although the clear effect that the ATR loss had on folliculogenesis, we observed a normal number of antral follicles in Seckel mouse ovaries. While we did not observe major defects in fetal oocytes, since previous reports failed to obtained viable oocytes after hormone-induced superovulation (Murga et al., 2009), we assume the oocytes from these antral follicles may not be fertilizable. Further investigations will have to elucidate this issue in more detail.

Finally, the data presented here have important implications for the use of ATR inhibitors as therapeutic drugs to treat certain kinds of cancers (Toledo et al., 2011; Weber and Ryan, 2015). Our investigations suggest that such treatments would block folliculogenesis progression in females, which could lead to infertility. Nonetheless, the fact that the primordial follicle pool is maintained in Seckel mouse, make us expect that the infertility originated by a treatment with ATR inhibitors would be transient and would not hamper the reproductive life of a patient after recovering from cancer. Thus, from a reproductive point of view, our results suggest that ATR inhibitors could be safe drugs in order to preserve female fertility after chemotherapy. Nevertheless, more studies need to address this issue in more detail, specially to study the possible effect of ATR inhibitors on dormant oocytes.

## Acknowledgements

We thank O. Fernández-Capetillo (CNIO, Spain) for providing us with the Seckel allele. This work was supported by the Ministerio de Ciencia e Innovación (BFU2010-18965, BFU2013-43965-P and BFU2016-80370-P, IR) and by the UAB-Aposta award to young investigators (APOSTA2011-03, IR).

---

**Supplementary figure 1. Follicle number per section is affected in adult Seckel mouse ovaries** Quantification of the number of follicles present at different stages in eight alternate ovarian sections from two ovaries of the indicated age and genotype. p values are from t test.

**Supplementary figure 2. Seckel mouse ovaries present and increased number of abnormal follicles** (**a**) Representative image of an abnormal follicle (AbF) exhibiting a large oocyte surrounded by one single layer of flattened granulosa cells. Note the presence of a regular primordial follicle (PF) next to it. Scale bar represents 50 μm. (**b**) Quantification of the number of abnormal follicles present in ovarian sections analyzed from the indicated genotypes. Columns and lines indicate the mean and standard deviation (SD).

**Supplementary figure 3. Seckel mouse oocytes occupy a larger area of the follicle than control ones** Indirect analysis of the number of follicular cells surrounding an oocyte. Each point represents the measure of the relative area occupied by an oocyte (blue circumference in the example image inset) in a particular follicle (green circumference) for the indicated genotypes at different stages of folliculogenesis. Horizontal black lines in the graph denote the means. Primary and small secondary follicles from Seckel ovaries present larger relative oocyte areas than control follicles, suggesting a reduction in the number of granulosa cells surrounding those oocytes.

**Supplementary Table 1.** Proportion of the area occupied by the oocyte in a follicle at different stages of folliculogenesis for the indicated genotypes.

|  | Primordial<br>follicles | Primary<br>follicles | Small<br>secondary<br>follicles | Pre-antral<br>secondary<br>follicles | Antral<br>follicles |
| --- | --- | --- | --- | --- | --- |
| ***Atr<sup>+/+</sup>*** | 0.7 ± 0.1, N<br>= 11 | 0.3 ± 0.1,<br>N = 62 | 0.3 ± 0.1, N<br>= 81 | 0.2 ± 0.1, N<br>= 67 | 0.04 ± 0.1,<br>N = 7 |
| ***Atr<sup>s/s</sup>*** | 0.7 ± 0.1,<br>N = 17 | 0.4 ± 0.2,<br>N = 67 | 0.4 ± 0.1, N<br>= 45 | 0.2 ± 0.1, N<br>= 68 | 0.5 ± 0.02, N<br>= 9 |

Mean ± standard deviation and total number of follicles analyzed (N) are shown.

## Materials and Methods

### Experimental animals

The *Atr* Seckel allele was generated previously (Murga et al., 2009). These mice were maintained in a C57BL/6 - 129/SV mixed genetic background. All the experiments were performed using at least two mutant animals and compared to littermate control mice. In those cases in which appropriate littermate controls were unavailable, control animals were obtained from other litters of the same cross and the same age. All the animals used for experimental procedures were sacrificed using CO2 euthanize methods. Experimental procedures performed in the present work conform to the protocol CEEAAH 1091 (DAAM6395) approved by the Ethics Committee for Animal Experimentation of the Universitat Autònoma de Barcelona and the Catalan Government. Previously reported methods were used to genotype the mice (Pacheco et al., 2018).

### Cytological methods

To obtained fetal oocyte spreads to study meiotic prophase, ovaries from 18 days *post partum* fetuses were harvested and digested in Collagenase in M2 medium for 30 minutes at 37^a^C, then treated with hypotonic buffer (30mM Tris-HCl pH=8.2, 50mM Sucrose, 17mM Sodium Citrate, 5mM EDTA, 0.5mM DTT, 1x Protease Inhibitory CocktaiI) for 30 minutes and oocytes were released in 100mM sucrose. Oocytes were transfered onto slides containing fixative (1% PFA, 5 mM Sodium Borate, 0.15% Triton X-100, 3 mM DTT, 1x PIC). After 2 hours, slides were washed with 4% Photoloflo, air-dried and stored at −80°C until use.

### Histological procedures

Ovarian samples were obtained from females at different ages depending on the experimental procedures. Juvenile and adult ovaries were fixed overnight in 4% paraformaldehyde in PBS or in Bouin’s solution (70% Saturated picric acid, 25% Formaldehyde, 5% Glacial acetic acid). Afterwards, samples were dehydrated, cleared and embedded in paraffin using standards procedures. 6μm thick sections were stained with PAS-hematoxylin. In order to detect and quantify apoptosis in ovarian sections, we performed TUNEL staining using the ApopTag Plus Fluorescein *In Situ* Apoptosis Detection kit (Millipore) following manufacturer’s instructions.

### EdU incorporation and assay

To measure follicular cells proliferation rate, we performed an intraperitoneal administration of 300 μl of the thymidine analog EdU (5-ethynyl-2’-deoxyuridine) (EdU 1 mg/ml in PBS) to female mice. 24h after administration, ovaries were harvested and fixed in 4% paraformaldehyde in PBS overnight. After two PBS washes, ovaries were incubated in a sucrose solution (30% sucrose in PBS) overnight at room temperature to cryoprotect the tissue. Samples were then embedded in OTC compound to form a block at −20°C. 10μm thick cryosections were obtained using a cryostat. Slides were permeabilized in 0.5% Triton X-100 in PBS and then blocked in 3% BSA in PBS. EdU detection was performed using a Click-It EdU Imagine Kit (Invitrogen) following manufacturer’s instructions. Afterwards, slides were immunostained to detect oocytes (see below).

### Immunofluorescence

Immunofluorescence was performed using standard methods described elsewhere (Roig et al., 2004). Primary antibodies used were: mouse anti SYCP3 (1:200, Abcam), rabbit anti SYCP3 (1:200, Abcam), mouse anti SYCP1 (1:400, Abcam), mouse anti γH2AX (1:200, EMD Millipore), mouse anti MLH1 (1:50, BD Bioscience), Rabbit anti DDX4 (1:100, Abcam).

### Microscopy, image processing and data analysis

PAS-Hematoxylin stained tissue sections were analyzed on a brightfield microscope Olympus CH2 and images were captured using Zeiss Axiophot Microscope and Olympus C5060 camera. For fluorescent sample analysis and image capturing a Zeiss Axioskop fluorescence microscope connected with a ProgRes Jenoptik camera was used. The image capture software ProgRes CapturePro was employed for image acquisition and image processing.

All microscopy images were processed and different fluorescent channels overlaid using Adobe Photoshop CS2. The Java-based image processing program ImageJ (imagej.nih.gov/ij/) was used to measure length and areas.

Data analysis and statistical significance inference were performed using the GraphPad InStat, GraphPad QuickCalcs (http://www.graphpad.com/quickcalcs/) and GraphPad Prism 5 softwares.

